# Bacterial competition mediated by siderophore production among the human nasal microbiota

**DOI:** 10.1101/432948

**Authors:** Reed M. Stubbendieck, Daniel S. May, Marc G. Chevrette, Mia I. Temkin, Evelyn Wendt-Pienkowski, Julian Cagnazzo, Caitlin M. Carlson, James E. Gern, Cameron R. Currie

## Abstract

Resources available in the human nasal cavity are limited. Therefore, to successfully colonize the nasal cavity, bacteria must compete for scarce nutrients. Competition may occur directly through interference (e.g., antibiotics) or indirectly by nutrient sequestration. To investigate the nature of nasal bacterial competition, we performed co-culture inhibition assays between nasal Actinobacteria and *Staphylococcus* spp. We found that *Staphylococcus epidermidis* isolates were sensitive to growth inhibition by Actinobacteria but *Staphylococcus aureus* isolates were resistant to inhibition. Among Actinobacteria, we observed that *Corynebacterium* spp. were variable in their ability to inhibit *S. epidermidis.* We sequenced the genomes of ten *Corynebacterium* spp. isolates, including three *Corynebacterium propinquum* that strongly inhibited *S. epidermidis* and seven other *Corynebacterium* spp. isolates that only weakly inhibited *S. epidermidis.* Using a comparative genomics approach, we found that the *C. propinquum* genomes were enriched in genes for iron acquisition and encoded a biosynthetic gene cluster (BGC) for siderophore production, absent in the non-inhibitory *Corynebacterium* spp. genomes. Using a chromeazurol S assay, we confirmed that *C. propinquum* produced siderophores. We demonstrated that iron supplementation rescued *S. epidermidis* from inhibition by *C. propinquum*, suggesting that inhibition was due to iron restriction through siderophore production. Using comparative metabolomics, we identified the siderophore produced by *C. propinquum* as dehydroxynocardamine. Finally, we confirmed that the dehydroxynocardamine BGC is expressed *in vivo* by analyzing human nasal metatranscriptomes from the NIH Human Microbiome Project.

Together, our results suggest that bacteria produce siderophores to compete for limited available iron in the nasal cavity and improve their fitness.

**IMPORTANCE:** Within the nasal cavity, interference competition through antimicrobial production is prevalent. For instance, nasal *Staphylococcus* spp. strains can inhibit the growth of other bacteria through the production of nonribosomal peptides and ribosomally synthesized and post-translationally modified peptides. In contrast, bacteria engaging in exploitation competition modify the external environment to prevent competitors from growing, usually by depleting access to essential nutrients. As the nasal cavity is a nutrient limited environment, we hypothesized that exploitation competition occurs in this system. We determined that *Corynebacterium propinquum* produces an iron-chelating siderophore and is able to use this molecule to sequester iron and inhibit the growth of *Staphylococcus epidermidis.* Further, we found that the genes required for siderophore production are expressed *in vivo.* Thus, though siderophore production by bacteria is often considered a virulence trait, our work indicates that bacteria may produce siderophores to compete for limited iron in the human nasal cavity.

## INTRODUCTION

Humans engage in symbioses with diverse sets of microbes at nearly every body site. Collectively, these microbes are referred to as the human microbiota. Members of the microbiota provide their human host with essential services. For example, within the gastrointestinal tract, mutualistic bacteria metabolize recalcitrant nutrients to release metabolites that are accessible to the human host (1, 2), synthesize many essential amino acids, cofactors and vitamins (3, 4), and provide defense against pathogens (5, 6). However, despite providing critical services to their hosts, nutrient acquisition is often a principle challenge for bacteria colonizing humans and other animals. In part, this is due to host sequestration of resources as a means to control bacterial growth (7).

To form and maintain associations with humans and other eukaryotes, bacteria have evolved mechanisms to derive nutrition from their hosts (8, 9). Both commensal and pathogenic bacteria colonizing the gastrointestinal tract consume host-derived compounds (10). For instance, *Bacteroides acidifaciens* and *Akkermansia muciniphila* cells have been shown to incorporate carbon and nitrogen from host proteins (11) and the pathogen *Salmonella enterica* serotype Typhimurium uses tetrathionate derived from thiosulphate produced by the host inflammatory response during infection as a terminal electron acceptor (12). In addition to consuming host-derived products, bacteria colonizing the gastrointestinal tract can also acquire nutrients from meals that their host consumes (10). However, outside of the gastrointestinal tract, bacteria have increasingly limited access to nutrients. *Cutibacterium* (*Propionibacterium*) *acnes* colonizing the sebaceous glands in human skin hydrolyzes sebum triglycerides to generate free fatty acids and glycerol (13, 14), which they may metabolize (15). Similarly, anaerobic oral bacteria that have spread to the lower respiratory tracts of cystic fibrosis patients may ferment host mucins to generate short chain fatty acids and amino acids to feed other bacteria, including *Pseudomonas aeruginosa* (16).

To colonize the upper airway, bacteria must attach to the epithelial surface, accommodate or evade the host’s immune system, and cope with decreased access to freely available nutrients when compared to bacteria colonizing the gastrointestinal tract (reviewed in 7). Within the human nasal cavity there are minute concentrations of free amino acids, carbohydrates, organic acids, and minerals (17–19). For comparison, nasal secretions contain ~65-fold lower glucose concentrations than the lumen of the small intestine (19, 20). Further, humans actively deplete the available glucose in the airway through polarized glucose importers in the membranes of airway epithelial cells (21). Similar to other environments such as the soil or the ocean, the bioavailability of iron in the nasopharynx is low (17, 18). As a means to acquire scarce iron from their environment, bacteria release iron-chelating molecules called siderophores that scavenge ferric iron and other minerals and are resorbed by their producers (22). However, human hosts have evolved countermeasures to circumvent bacterial siderophores. For instance, bacterial colonization of the nasopharynx triggers neutrophils to produce lipocalin-2, a protein that binds to enterobactin-type siderophores and prevents their uptake by bacteria (23, 24). Finally, in addition to experiencing nutrient and mineral limitation, bacteria that colonize the nasal cavity are exposed to oxygen stress, which may be either abiotic or produced by the action of host immune cells (25, 26). Therefore the human nasal cavity is a low resource and high stress environment, requiring bacteria in this environment to engage in competitive interactions for survival (27, 28).

Competition is split into two modes called interference and exploitation. Bacteria engaging in exploitation competition compete by preventing their competitors from accessing resources, either by rapidly consuming or sequestering them. In contrast, bacteria using interference competition use toxic effectors to directly inhibit their competitors (29). Bacteria from the human nasal cavity are well known to use specialized (secondary) metabolites with antimicrobial properties to engage in interference competition. As examples, *Staphylococcus lugdunensis* produces a thiazolidine-containing cyclic peptide called lugdunin that inhibits the growth of *Staphylococcus aureus* (30). *Corynebacterium accolens* secretes a lipase that cleaves human nasal triacylglycerols to produce antimicrobial free fatty acids that inhibit the growth of *Streptococcus pneumoniae* (31). Further, under conditions of iron-limitation and hydrogen peroxide induced oxidative stress, which reflect conditions experienced by bacteria colonizing the nasal cavity, *Staphylococcus epidermidis* IVK45 increases production of an antimicrobial peptide called nukacin IVK45 (32). Despite these examples of interference competition and the low resource environment of the nasal cavity, mechanisms of exploitation competition are not well studied among the human nasal microbiota (33).

In this study, we investigated bacterial competition between Actinobacteria and *Staphylococcus*, which are among the most abundant members of the human nasal microbiota (33, 34). We found that *Corynebacterium* (Phylum: Actinobacteria) vary in their ability to inhibit the growth of *S. epidermidis.* Using a comparative genomics approach, we identified a gene cluster for siderophore biosynthesis that is present in the genomes of *Corynebacterium propinquum* strains that more strongly inhibit *S. epidermidis.* We confirmed the siderophore production and demonstrate that iron sequestration as the mechanism of *S. epidermidis* inhibition. We identified the siderophore as dehydroxynocardamine. Finally, we detected expression of the dehydroxynocardamine biosynthetic gene cluster (BGC) in metatranscriptomic reads from the human nasal cavity. Together, the data suggest that members of the human nasal microbiota engage in exploitation competition for limiting iron, and may influence the composition of the human nasal microbiota *in vivo.*

## MATERIALS AND METHODS

### Nasal lavage specimen collection

The nasal lavage samples that we used in this study were banked as part of the Childhood Origins of ASThma (COAST) study (35). Informed consent was obtained from the parents, and the Human Subjects Committee at the University of Wisconsin-Madison approved the study (IRB approval number H-2013–1044). Briefly, lavage samples were collected by spraying Deep Sea Nasal Spray (Major^®^) into one of the participant’s nostrils. To collect the sample, the participant was then instructed to blow their nose into a plastic bag. Subsequently, the samples were stored at 4 °C. Approximately 3 mL of either phosphate-buffered saline or Amies transport media (Copan Diagnostics) was added to each sample before storage at −80 °C.

### Strain isolation and maintenance

For general strain propagation, we used brain heart infusion (BHI) (DOT Scientific). All plates were solidified using 1.5% agar (VWR). For iron supplementation experiments, we added 200 μM FeCl_3_ to the BHI agar after autoclaving and cooling the media to 55 °C before pouring the plates. The strains of bacteria used in this study are listed in **Table S1**. To culture bacteria from the lavage samples, we spread 100 μL of each of the thawed samples onto BHI plates, and incubated the plates aerobically at 37 °C for 1 week. We selected ≥2 colonies of each distinct morphotype per plate and passaged the isolates aerobically on BHI plates at 37 °C until we obtained pure cultures. All bacterial isolates were cryopreserved at −80 °C in 25% glycerol.

### Bacterial isolate identification

To identify bacterial isolates from the nasal lavage samples, we sequenced the 16S *rRNA* gene. We used colony PCR with the universal 27F (5’-AGAGTTTGATCMTGGCTCAG-3’) and 1492R (5’-CGGTTACCTTGTTACGACTT-3’) primers to amplify the 16S *rRNA* gene (36). We sequenced PCR products using the Sanger method at the University of Wisconsin-Madison Biotechnology Center. We identified isolates to the genus level using the Ribosomal Database Project Classifier (37). To distinguish between *S. aureus* and *S. epidermidis* isolates, we used BLAST searches against the NCBI 16S ribosomal RNA sequence database with a threshold sequence identity of 99%. We corroborated the identity of *Staphylococcus* spp. isolates using colony pigmentation and hemolysis phenotypes (38–40).

### 16S rRNA gene phylogenetic analysis

We aligned the 16S *rRNA* gene sequences using SINA v1.2.11 (41) through the Silva webserver (42) and discarded unaligned bases at the ends of the sequences. For phylogenetic analysis, we imported the alignments into MEGA7 (43). We inferred phylogenetic trees using the maximum likelihood method based on the general time reversible model. All nucleotide positions containing gaps were removed in the final analysis.

### Co-culture plate inhibition assays

We used co-culture plate inhibition assays to assess the activity of nasal Actinobacteria isolates against nasal *Staphylococcus.* We spread Actinobacteria from saturated overnight cultures in BHI broth over one-half of a well (d = 2.4 cm) containing 3 mL of BHI agar on a 12-well plate (Greiner Bio-One). We incubated the plates for one week at 37 °C. Subsequently, we spotted ~3 μL of a ten-fold diluted saturated overnight culture of each *Staphylococcus* spp. isolate in BHI adjacent to the Actinobacteria and returned the plates to the 37 °C incubator. After 1 week of co-culture, we removed the plates, scanned them, and scored the inhibition for each interaction pair. All interactions were tested with ≥2 replicates with consistent results. An inhibition score of 0 indicated no inhibition, a score of 1 indicated weak inhibition, and a score of 2 indicated strong inhibition (**Fig. 1A**). We determined significance using the chi-square test.

**Figure 1.**
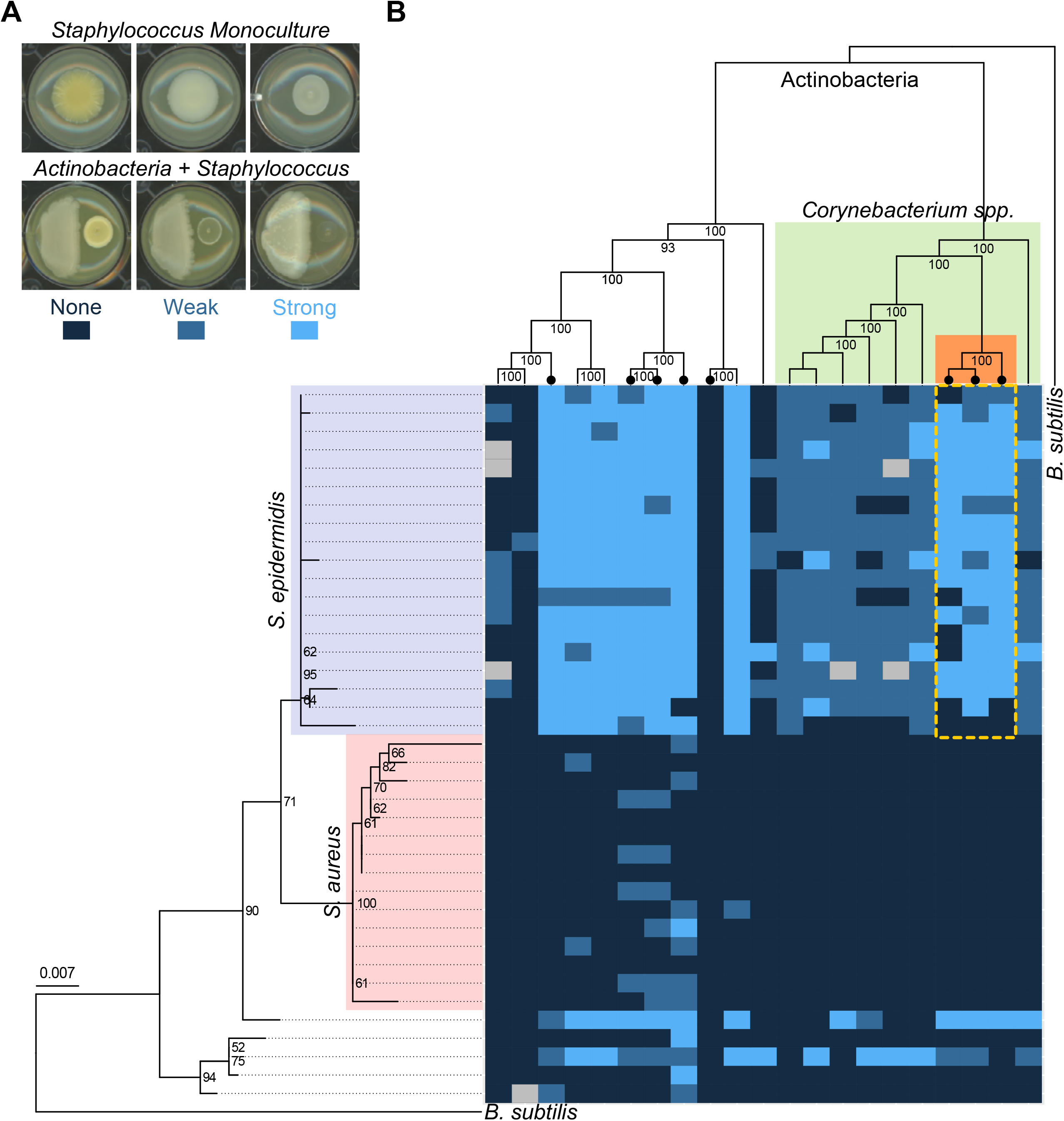
Inhibition of nasal *Staphylococcus* spp. by nasal Actinobacteria. **(A)** Representative images of monocultures of *Staphylococcus* and co-cultures of Actinobacteria (left) with *Staphylococcus* spp. isolates (right). The inhibition scoring system is depicted below the co-culture images. **(B)** 21 nasal Actinobacteria isolates (horizontal) were monocultured on BHI agar wells for 1 week before 39 *Staphylococcus* spp. isolates (vertical) were spotted adjacent to the Actinobacteria colony. The colonies were cultured together for one week before inhibition of the *Staphylococcus* spp. colony was scored. The heat map displays the inhibition scores of each *Staphylococcus* spp. isolate when paired with the corresponding Actinobacteria isolate. Each interaction was technically replicated at least twice. The gray cells indicate interactions where replicates were in disagreement or the Actinobacteria colony overgrew the well before *Staphylococcus* inoculation. (*left*) Phylogenetic tree of *Staphylococcus* spp. isolates built from the 16S rRNA gene. (*top*) Core-genome phylogenetic tree of Actinobacteria built from 93 conserved, single copy genes. Both phylogenies are rooted on *B. subtilis* 168 and nodes with ≥50% bootstrap support are indicated. The dashed yellow lines highlight strong inhibition of *S. epidermidis* by one clade of *Corynebacterium* spp. Actinobacteria taxa marked with the • symbol encode a siderophore BGC within their genome.

### Whole genome sequencing and assembly

We cultured bacterial strains for whole genome sequencing in 3 mL of BHI broth supplemented with 0.5% glycine overnight at 37 °C. We harvested the cells by centrifuging the cultures at 21,330×g for 5 min. The cell pellets were washed with 10.3% sucrose and resuspended in 450 μL of lysozyme solution [3 mg/mL lysozyme (Sigma), 0.3 M sucrose, 25 mM Tris-HCl (pH 8), 25 mM EDTA (pH 8)] for 30 min at 37 °C. Subsequently, 13 μL of 20 mg/mL Proteinase K (Thermo Fisher) was added to each sample, with additional incubation at 42 °C for 15 min. Cells were lysed by adding 250 μL of 2% SDS and rocking for 15 min. DNA was purified using standard phenol-chloroform extraction and precipitated with 3 M sodium acetate and isopropanol. We visually assessed the quality of genomic DNA preparations by running samples on 0.5% TBE gels. Genomic libraries for Illumina MiSeq 2 × 150 bp paired-end sequencing were prepared and sequenced by the University of Wisconsin-Madison Biotechnology Center. Raw reads were corrected with MUSKET v1.1 (44) and paired ends were merged with FLASH v1.2.7 (45). Reads were assembled into draft genome sequences with SPAdes v3.11.0 (46).

The genome sequences of Actinobacteria generated during this study have been deposited in the National Center for Biotechnology Information database under BioProject accession PRJNA492917.

### Core-genome phylogeny

To construct the core-genome phylogenetic tree of nasal Actinobacteria, we used the core species tree pipeline (https://github.com/chevrm/core_species_tree) developed in our laboratory (47). Briefly, we called genes in each genome using prodigal v2.6.0 (48) and used profile Hidden Markov Models in HMMER v3.1b2 (49) to search each genome for 93 full length TIGRFAM amino acid sequences in the “core bacterial protein” set (GenProp0799). Each protein family was aligned using MAFFT v7.245 (50) and converted to codon alignments. We used RAxML v8.1.24 (51) to generate phylogenetic trees for each of the 93 codon alignments under the GTRgamma substitution model with 100 bootstraps. To generate the species phylogenetic tree, we used ASTRAL-II (52) from the individual phylogenetic trees with 100 bootstraps. We used FigTree v1.4.3 (http://tree.bio.ed.ac.uk/software/figtree/) to root the phylogeny on *B. subtilis* 168 and display the branch length based on proportional length to the root.

### Comparative genomics

We annotated the assembled genomes using prokka v1.13 (53). For comparative genomics, we identified orthologs from the nucleotide sequences of each protein-coding open reading frame (ORF) with OMA v2.2.0 (54). We applied Euclidean hierarchical clustering to the ortholog presence/absence matrix generated by OMA using R. To annotate orthologues with Kyoto Encyclopedia of Genes and Genomes (KEGG) orthology (KO) terms, we used DIAMOND v0.9.21.122 (55) to query one sequence of each ortholog group against a custom database of KEGG annotated proteins sequences with an E-value threshold cutoff of 1.0e-10. Each ORF was assigned the KO terms from the top BLAST hit. For KEGG KO term enrichment analysis, we used the GSEABase package v1.42.0 (56) and GOstats v1.7.4 (57) in R and performed a hypergeometric test with a *p*-value cutoff of 0.05.

### Identification of BGCs

To identify BGCs we used antiSMASH 4.0.2 (58). Proteins from all identified BGCs were aligned via an all versus all DIAMOND (query cover >75, percent identity >60). BGCs that shared over 75% of proteins were called the same family. BGC families correlated to inhibition were identified through presence in the high-inhibition strains and absence in the weak-inhibition strains. Proteins from all identified BGCs were combined with all proteins from MIBiG v1.3 (59) and subsequently aligned with DIAMOND with the same parameters as listed before to identify biosynthetic gene similarities to previously described BGCs.

### Chrome azurol S assay

We used the overlay chrome azurol S (CAS) assay to test for siderophore production as previously described (60). We cultured *Corynebacterium* spp. overnight in 3 mL BHI at 37 °C. We diluted the overnight cultures to OD_600_ = 2 and spotted 5 μL on to 10 mL BHI agar plates (d = 5 cm). We incubated the plates at 37 °C for 4 days to ensure robust *Corynebacterium* spp. growth. Subsequently, we overlaid each plate with 6 mL of CAS reagent overlay [100 μM CAS, 200 μM hexadecyltrimethylammonium bromide, 10 μM FeCl_3_·6 H_2_O, 10 mM PIPES (pH 6.8), 1% agarose] (60, 61) and incubated the plates at ambient temperature in the dark overnight before scanning plates for color change from blue to yellow, indicating production of a siderophore (61).

### Siderophore identification and mass spectral molecular networking

To identify the siderophore, we used a mass spectrometry guided approach. We spotted 10 μL of either *Corynebacterium genitalium* HSID17231 or *Corynebacterium propinquum* HSID18034 overnight cultures onto the center of 25 mL BHI agar plates (d = 8.5 cm). After incubating the plates as above, we removed agar cores (d = 0.6 cm) from near the *Corynebacterium* spp. colony using the wide end of a P1000 pipette tip. As above, we verified siderophore activity by directly placing duplicate agar cores onto CAS assay plates and observing the color change from blue to yellow. We washed the agar cores in 2 mL of 50% methanol and 2 mL of 100% methanol, which we combined and dried with a gentle air stream under reduced pressure.

Data-dependent LC/MS^2^ was run on a Thermo Scientific Q Exactive Orbitrap by the University of Wisconsin Analytical Instrumentation Center of the School of Pharmacy. The liquid chromatography method was run on a Phenomenex XB C_18_, 2.1 × 100 mm, 2.6 μm particle size column with solvents A (water with 0.1% formic acid) and B (acetonitrile with 0.1% formic acid). A gradient of 5% B for half a minute to 30% B over 16 minutes, then 97% B for two minutes with a flow rate of 0.35 mL/min was used to separate the metabolites. We exported the files in mzXML format and uploaded them to Global Natural Products Social Molecular Networking (GNPS) (62). We generated a molecular network using the online workflow at GNPS. The data was filtered by removing all MS/MS peaks within ± 17 Da of the precursor *m/z.* The MS^2^ spectra were window filtered by choosing only the top 6 peaks in the ± 50 Da window throughout the spectrum. The data was then clustered with MS-Cluster with a parent mass tolerance of 2.0 Da and a MS^2^ fragment ion tolerance of 0.5 Da to create consensus spectra. Consensus spectra that contained less than 2 spectra were discarded. A network was then created where edges were filtered to have a cosine score above 0.6 and more than 3 matched peaks. Edges between two nodes were kept in the network if each of the nodes appeared in the respective top 10 most similar nodes. The spectra in the network were then searched against GNPS’ spectral libraries. The library spectra were filtered in the same manner as the input data. All matches kept between network spectra and library spectra were required to have a score above 0.6 and at least 3 matched peaks. Analog search was enabled against the library with a maximum mass shift of 100.0 Da.

### Metatranscriptome analysis for dehydroxynocardamine BGC expression

As part of the National Institutes of Health (NIH) Integrative Human Microbiome Project (iHMP), RNA from the anterior nares of 16 prediabetic subjects was isolated and depleted of 16S and 23S rRNAs. The RNA was converted to cDNA with random primers and paired end sequenced using an Illumina platform (63). We downloaded the raw reads from the iHMP using the hmp_client (https://github.com/ihmpdcc/hmp_client) from the Human Microbiome Project Data Portal (https://portal.hmpdacc.org). We removed Illumina adapter sequences, low quality reads, and unpaired reads using trimmomatic v0.36 (64) with the following parameter: SLIDINGWINDOW:5:20. RNA was isolated from each of the 16 subjects between 1 and 11 times for a total of 95 nasal metatranscriptomes (mean = 5.9/subject, median = 6.5/subject). For our analyses, we treated each individual metatranscriptome as a single sample. To determine expression of the dehydroxynocardamine BGC, we used kallisto version 0.44.0 (65) and pseudoaligned each of the metatranscriptomes onto the indexed open reading frames (ORFs) onto the *Corynebacterium propinquum* HSID18034 genome. We extracted the expression data for the combined dehydroxynocardamine BGC (*dnoA-G*) and four housekeeping genes: *rpoB, gyrB, sigA*, and *rpsL.* We removed genes with <1 estimated count from the dataset for a total of 148 expression counts (18 siderophore BGC, 130 housekeeping genes).

### Plot generation and data visualization

We generated all plots in R v3.5.0 (66) using the ggplot2 package v2.2.1 (67). We visualized phylogenetic trees in R using the ggtree v1.12.0 package (68).

## RESULTS

### Interaction assays reveal variation in staphylococcal inhibition by nasal Actinobacteria

The scarcity of nutrients and minerals in the human nasal cavity results in a stressful environment where the microbiota must compete to survive. To pursue mechanisms of competition that occur among the nasal microbiota, we chose to use a culture-based approach and investigate interactions between Actinobacteria and *Staphylococcus* spp., which are major bacterial colonizers of the human nasal cavity. For this study, we isolated bacteria from frozen nasal lavage samples donated by healthy children as part of the COAST study (35, 69), which we identified by colony morphology, pigmentation, hemolysis, and 16S rRNA gene sequence (see Methods).

To identify differences in patterns among interactions between Actinobacteria and *Staphylococcus* spp., we assessed the ability of a subset of Actinobacteria isolates to inhibit the growth of a subset of *Staphylococcus* spp. isolates. In total, we tested 21 Actinobacteria isolates (10 *Corynebacterium*, 1 *Curtobacterium*, 1 *Dermabacter*, 3 *Kocuria*, 1 *Microbacterium*, 3 *Micrococcus*, and 2 *Rothia*) against 39 *Staphylococcus* spp. isolates (15 *S. aureus*, 19 *S. epidermidis*, and 5 *S. warneri/pasteuri)* (**Table S1**) for a total of 812 pairwise combinations with ≥2 replicates. In the case of 7 pairwise combinations, the replicates were not in agreement or the Actinobacteria isolate overgrew its well and prohibited inoculation of the *Staphylococcus* spp. isolate. We removed these 7 points from further analysis. We scored inhibition by visual inspection of the *Staphylococcus* spp. colonies after 1 week of co-incubation. Strong inhibition corresponded to colonies with severe growth defects or total growth inhibition, whereas weak inhibition corresponded to colonies with moderate inhibition resulting in diminished growth. *Staphylococcus* spp. colonies that were indistinguishable to a monoculture control were considered uninhibited (**Fig. 1A**). To visualize patterns in the inhibition assays, we plotted the inhibition scores as a heat map that was clustered based on the phylogenetic relationships of the Actinobacteria and *Staphylococcus* spp. isolates on the horizontal and vertical dimensions, respectively. The *Staphylococcus* spp. phylogenetic tree was built from 16S *rRNA* gene sequence alignments, whereas the Actinobacteria phylogenetic species tree was built from alignments of 93 single copy core bacterial genes that were extracted after whole genome sequencing of the Actinobacteria isolates (**Fig. 1B**).

By clustering the inhibition scores with respect to phylogeny, we immediately observed significant differences in the inhibition patterns between *S. aureus* and *S. epidermidis.* Broadly, most *S. epidermidis* isolates were inhibited by nasal Actinobacteria but the *S. aureus* isolates were insensitive [*χ*^2^ = 371.73, *df* = 2, *p* < 2.2e-16] (**Fig. 1B**). However, the inhibition pattern was not solely dominated by the identity of the *Staphylococcus* species; we observed variation among *Corynebacterium* spp. isolates to inhibit *S. epidermidis.* In particular, we noticed 3 *Corynebacterium* spp. isolates forming a monophyletic clade (highlighted in yellow in **Fig. 1B**) that more strongly inhibited the growth of *S. epidermidis* than the other *Corynebacterium* spp. isolates [*χ*^2^ = 89.74, *df* = 2, *p* < 2.2e-16] (**Fig. 1B**). Given the limited variation in staphylococcal inhibition by other genera of Actinobacteria, we focused on *Corynebacterium* spp. to determine underlying factors responsible for the differences in inhibition patterns.

### *Corynebacterium propinquum* strains strongly inhibit *S. epidermidis*

To identify the *Corynebacterium* strains, we constructed a core genome phylogeny from the ten *Corynebacterium* spp. that we isolated and 45 human-associated strains whose sequences were available from GenBank. All three of the strong inhibitors of *S. epidermidis* were mostly closely related to *Corynebacterium propinquum*, while the weak inhibitors of *S. epidermidis* were either closely related to *Corynebacterium pseudodiphtheriticum* (5 strains) or *Corynebacterium genitalium* (1 strain) (**Fig. S1**). For clarity, we will refer to the *Corynebacterium* strong inhibitors of *S. epidermidis* as *C. propinquum* and the other isolates as *C. pseudodiphtheriticum* or *C. genitalium.*

### Iron acquisition by *Corynebacterium* spp. is related to increased *S. epidermidis* inhibition

To identify differences among the *Corynebacterium* spp. strains that corresponded to differential *S. epidermidis* inhibition, we took a comparative genomics approach. We generated draft genome sequences for each of the ten *Corynebacterium* spp. isolates. Using the OMA algorithm, we predicted orthologs across the ten genomes. In total, we predicted 2702 ortholog groups, with 1198 groups (44% total) universally shared across the ten *Corynebacterium* spp. strains and 1504 ortholog groups (56% total) that were absent in at least one strain. In particular, the gene content in *Corynebacterium genitalium* HSID17239 was more divergent than the other isolates (**Fig. 2A, top row**). Indeed, when *C. genitalium* HSID17239 was excluded from the analysis, 1801 of the 2702 (67% total) ortholog groups were shared universally across the remaining nine strains. To observe differences in the patterns of gene content within the *Corynebacterium* spp. genomes, we applied hierarchical clustering to the presence-absence matrix of the 2702 ortholog groups. There were 337 orthologs (12% total) that were uniquely shared among *C. propinquum* (**Fig. 2A**). Though 223 of the orthologues were annotated as encoding hypothetical or putative proteins, a manual investigation of the 114 annotated orthologues revealed that 31 (27.2% of subset) were annotated with functions related to iron acquisition and siderophore biosynthesis. This result represents a 6.4-fold enrichment over the 4.3% abundance (68/1596 orthologs) of the corresponding terms in all ten *Corynebacterium* genomes (hypergeometric test: expected = 4.9%, *p* = 1.3e-19). As a second test, we performed an enrichment analysis for KEGG orthology (KO) terms. We annotated 169 of the 337 orthologues (50% of subset) with 117 KO terms. When compared to the total set of 357 KO terms, the 169 orthologues were enriched in 12 KO terms. In particular, we noted that among the KO terms, 2 KO terms were associated with iron acquisition and siderophore biosynthesis: 1) iron complex transport system permeases and 2) bifunctional isochorismate lyase/aryl carrier proteins (**Fig. 2B**).

**Figure 2.**
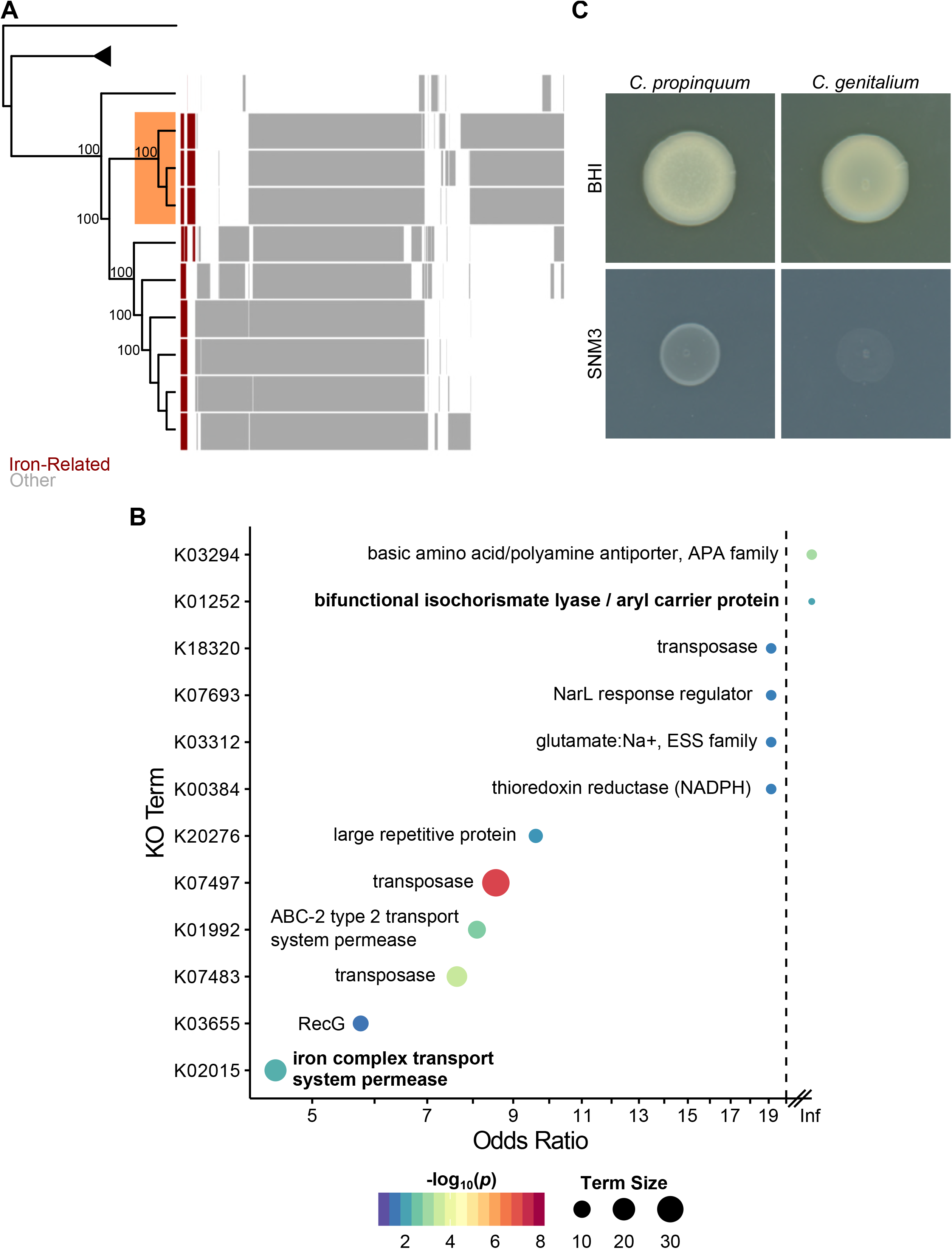
Comparative genomics of nasal *Corynebacterium* spp. **(A)** (*left*) The core-genome phylogeny of nasal *Corynebacterium* from **Fig. 1**. The node corresponding to other Actinobacteria is collapsed and represented by a triangle. The *C. propinquum* clade is shaded in orange. (*right*) Clustered presence-absence matrix of 1504 orthologues predicted across the 10 *Corynebacterium* spp. genomes. The 1198 orthologues that were conversed across all 10 genomes were excluded. Orthologs encoding functions pertaining to iron metabolism, iron transport, or siderophore biosynthesis are depicted in red. **(B)** KO term enrichment analysis of the orthologues from highlighted box in panel **A**, relative to the other orthologues. The odds ratios of all 12 significantly enriched (*p* < 0.05) KO terms are plotted. Note, an odds ratios of infinity (Inf) correspond to KO terms where all orthologues that were annotated with the KO term were present in the enrichment set. KO terms in bold are related to iron transport or siderophore biosynthesis. The color of each point indicates significance level and the size of the point indicates how many of the orthologues were annotated with the KO term. **(C)** Growth assay of *Corynebacterium* spp. strains that either weakly (*C. genitalium*) or strongly (*C. propinquum*) inhibited *S. epidermidis* on BHI and iron-limited SNM3.

The results of each set of enrichment analyses led us to hypothesize that *Corynebacterium* spp. that weakly inhibit *S. epidermidis* would have growth defects under conditions of iron limitation. To directly test for iron-related phenotypic differences between the two groups of *Corynebacterium*, we cultured strains on standard BHI media and on synthetic nasal medium 3 (SNM3), a medium that is formulated to replicate the low concentrations of nutrients and mineral content found in human nasal secretions (19). We observed no difference in the growth of *Corynebacterium* spp. strains on standard BHI media, but found that *C. genitalium* HSID17239 had a growth defect when cultured on SNM3 and was unable to form a colony (**Fig. 2C**). Together, these results demonstrate that different strains of *Corynebacterium* spp. isolated from the human nasal cavity possess differential ability to survive under iron-limited conditions, which correlates with the ability to inhibit the growth of *S. epidermidis.*

### *Corynebacterium propinquum* strains produce siderophores

The results of the gene and KO term enrichment analyses and iron-limitation phenotype assay (**Fig. 2**) led us to hypothesize that the observed variation in inhibition of *S. epidermidis* by *Corynebacterium* spp. (**Fig. 1B**) may be due to siderophore production. To directly detect siderophore production we used the CAS assay, which detects siderophore production by a color change from blue to yellow as siderophores remove iron from the CAS dye (61). Consistent with our previous results, we confirmed siderophore production by all three *C. propinquum* strains, whereas none of the other seven *Corynebacterium* spp. produced detectable siderophore activity (**Fig. 3A**).

**Figure 3.**
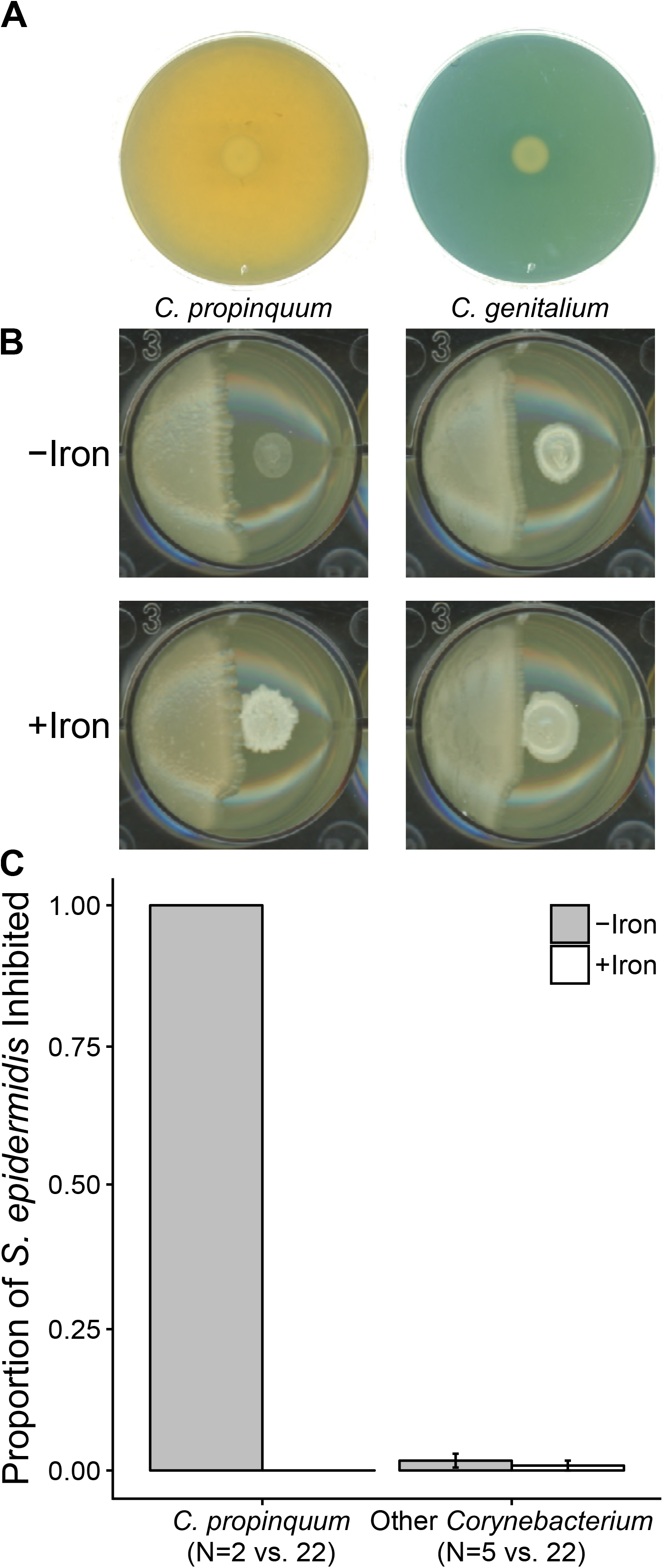
Siderophore production by *C. propinquum* inhibits *S. epidermidis.* **(A)** CAS assay results from *C. propinquum* HSID18034 (left) and *C. genitalium* HSID17239 (right), which strongly and weakly inhibit *S. epidermidis*, respectively. A shift in the color of the CAS overlay from blue to yellow indicates siderophore production. **(B)** Co-culture plate inhibition between the strains in **A** and *S. epidermidis* on BHI (−Iron) and BHI supplemented with 200 FeCl_3_ (−Iron). **(C)** Co-culture plate inhibition assays were performed between *C. propinquum* strains (HSID18034 and HSID18036) and five *Corynebacterium* spp. non-siderophore-producers (HSID17231, HSID17239, HSID17260, HSID17564, HSID17575) against 22 strains of *S. epidermidis* on BHI (−Iron) and BHI supplemented with 200 μM FeCl_3_ (+Iron). Each pairing was duplicated with consistent results. The error bars represent the standard error of the mean. The photographs in **A** and **B** are representative of duplicate samples.

### Iron supplementation rescues *S. epidermidis* from siderophore-mediated inhibition

The results from the CAS assay (**Fig. 3A**), indicated that siderophore production is correlated with strong inhibition of *S. epidermidis.* To directly test if iron sequestration by *C. propinquum* was responsible for *S. epidermidis* inhibition, we repeated the co-culture plate inhibition assays on BHI and on BHI supplemented with 200 μM FeCl_3_. We found that iron supplementation rescued *S. epidermidis* from inhibition by *Corynebacterium propinquum* (**Fig. 3B**). We performed the assay with two *C. propinquum* siderophore producers and five non-siderophore producers (1 strain of *C. genitalium* and 4 strains of *C. pseudodiphtheriticum)* against 22 strains of *S. epidermidis* isolated from different nasal specimens. On normal BHI medium, *C. propinquum* universally inhibited isolates of *S. epidermidis* and the inhibition was completely reversed when the medium was supplemented with FeCl_3_ (**Fig. 3B-C**), whereas under the conditions of this assay, there was little-to-no inhibition of *S. epidermidis* by either *C. genitalium* or *C. pseudodiphtheriticum* under either culture condition (**Fig. 3C**) [*χ*^2^ = 291.16, *df* = 3, *p* < 2.2e-16]. This result indicates that inhibition of *S. epidermidis in vitro* by *C. propinquum* is due to iron sequestration.

### The *C. propinquum-produced* siderophore is dehydroxynocardamine

As a first step to identify the siderophore produced by *C. propinquum*, we used antiSMASH to BGCs for siderophore biosynthesis in the *Corynebacterium* spp. genomes that we sequenced for this study. We identified a single 12553 bp siderophore BGC that was present within all three *C. propinquum* genomes but absent in the other seven *Corynebacterium* spp. genomes. This siderophore BGC was composed of seven open reading frames (ORFs), which were annotated with functions consistent with iron acquisition and siderophore biosynthesis (70–72) (**Fig. 4A**). Specifically, in addition to transport-associated genes, this BGC encodes a PLP-dependent decarboxylase, an L-lysine N6-monooxygenase, and a siderophore synthetase that contains acyl-CoA N-acyltransferase (IPR016181), aerobactin siderophore biosynthesis, IucA/IucC, N-terminal (IPR007310), and ferric iron reductase FhuF (IPR022770) domains. Though there were no experimentally characterized BGCs that were identical to the *C. propinquum* siderophore BGC, the order of the biosynthetic genes was identical to the desferrioxamine E *dfo* BGC from *Pantoea agglomerans* strain B025670 (MIBiG BGC0001572). The biosynthesis enzymes encoded by both siderophore BGCs are highly similar (**Table 1**) and both synthetases belong to the type C’ nonribosomal peptide synthetase-independent family (71, 72), which indicated that the siderophore produced by *C. propinquum* is likely macrocyclic.

**Figure 4.**
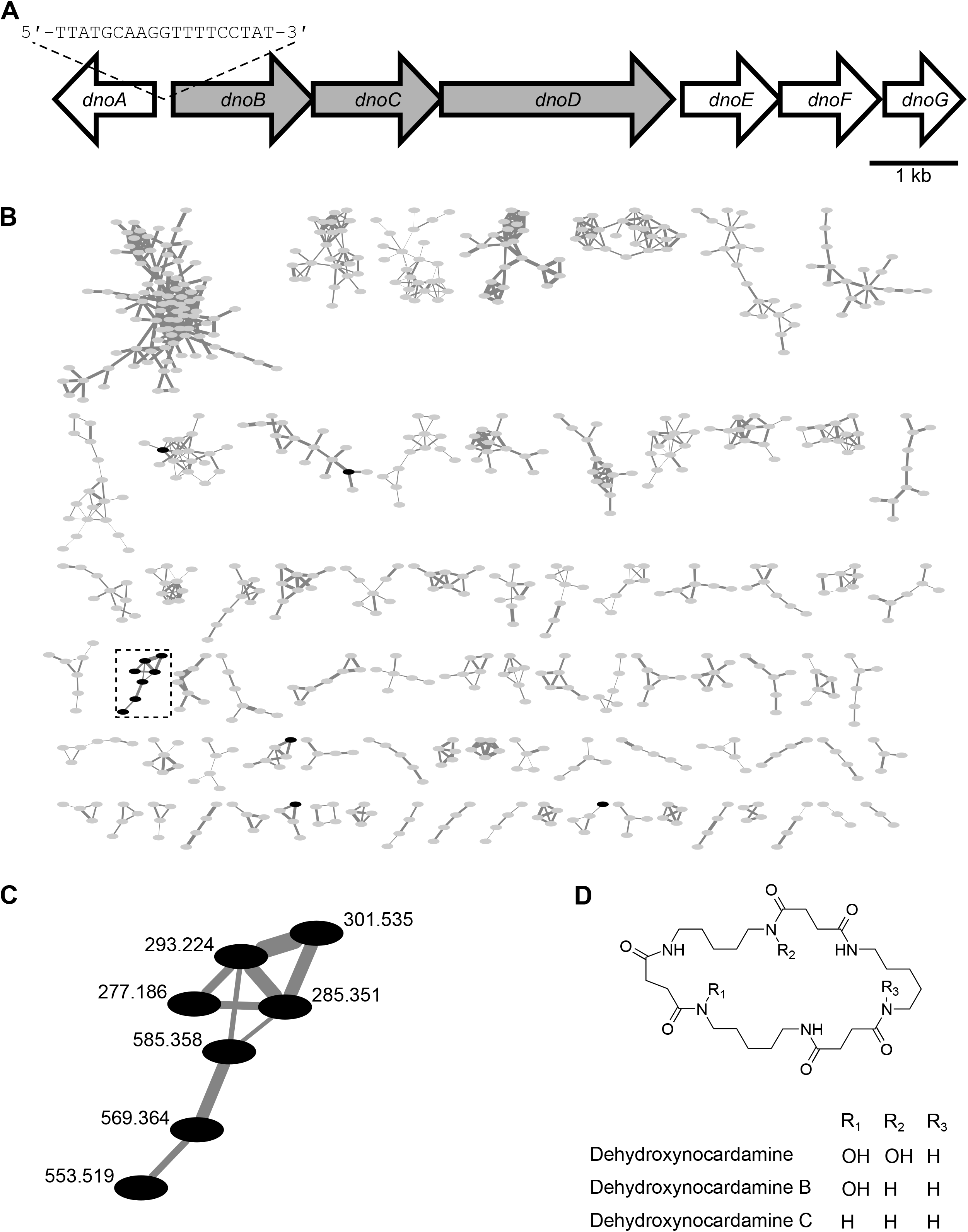
*Corynebacterium propinquum* produces the siderophore dehydroxynocardamine. **(A)** The dehydroxynocardamine BGC from *C. propinquum.* ORFs encoding biosynthetic and transport-related functions are filled with gray and white, respectively. See **Table 1** for specific ORF annotation. The DtxR sequence motif is shown upstream of *dnoB.* **(B)** Subset of the molecular network of *C. propinquum* and *C. genitalium* agar core extracts. Grey nodes are metabolites shared by both strains and black nodes are metabolites unique to *C. propinquum.* There were no detected metabolites that were unique to *C. genitalium. The* edges are weighted to the cosine score between the two features. A single cluster unique to *C. propinquum* is outlined in a dashed box. **See Fig. S2** for the full molecular network **(C)** Zoomed in view of the single cluster unique to *C. propinquum* with the nodes labeled by their corresponding *m/z.* This cluster contains single and double charged states of dehydroxynocardamine, dehydroxynocardamine B, and dehydroxynocardamine C. **(D)** Structures of dehydroxynocardamine, dehydroxynocardamine B, and dehydroxynocardamine C.

**Table 1.** *Corynebacterium propinquum* siderophore BGC annotation.

| ORF | Size (nt) | KO | Annotation | Desferrioxamine E BGC Homologue (E-Value*) |
| --- | --- | --- | --- | --- |
| *dnoA* | 1074 | 02016 | Iron-siderophore ABC transporter substrate-binding protein | n/a |
| *dnoB* | 1539 | 13745 | PLP-dependent decarboxylase | *dfoJ* (4e-116) |
| *dnoC* | 1152 | 03897 | L-lysine N6-monooxygenase | *dfoA* (1e-119) |
| *dnoD* | 2553 | n/a | siderophore biosynthetic enzyme | *dfoC* (2e-153) |
| *dnoE* | 1062 | 02015 | Ferric siderophore ABC transporter permease | n/a |
| *donF* | 1110 | 02015 | Siderophore ABC transport system permease | n/a |
| *dnoG* | 840 | 02013 | Siderophore transport system ATP-binding protein | n/a |

\*E-value from blastp protein sequence alignment.

To identify the siderophore produced by *C. propinquum*, we used comparative mass spectrometry-based metabolomics. We cultured *C. propinquum* HSID18034 and *C. genitalium* HSID17239 on BHI agar plates. We sampled agar cores near the bacterial colonies, which we extracted using methanol, and analyzed using data dependent liquid chromatography-tandem mass spectrometry. To visualize differences in the metabolomic profiles of the two strains, we used mass spectral molecular networking (73). Surprisingly, in a network containing 222 clusters and 1531 singletons (**Fig. 4B**), we observed only a single cluster that was unique to *C. propinquum* HSID18034 (**Fig. 4C**). The central node in this cluster corresponded to *m/z* 585.358 [M^+^H]^+^ (C_27_H_48_N_6_O_8_, Δppm = 3.44). The molecular weight and tandem mass spectrometry fragmentation pattern matched that of the siderophore dehydroxynocardamine (*m/z* 585.361 [M^+^H]^+^, C_27_H_49_N_6_O_8_), which is also called terragine E (**Fig. 4D**). This siderophore has been previously reported from marine sponge-associated *Streptomyces* (74) and produced by heterologous expression of soil DNA libraries in *Streptomyces lividans* (75), but has not been reported from *Corynebacterium* or any other human-associated bacteria. The other nodes in the cluster correspond to two dehydroxynocardamine analogs with protonated ions of *m/z* 569.364 [M^+^H]^+^ and 553.368 [M^+^H]^+^, as well as the doubly charged states of each molecule (**Fig. 4C**). The high similarity in fragmentation patterns as well as the differences of *m/z* −16 and −32 suggested these compounds contained one and two fewer N-hydroxy groups than dehydroxynocardamine (**Figs. 4D, S3**). Based on the product, we chose to name the *C. propinquum* siderophore BGC *dno* for <u>d</u>ehydroxy<u>no</u>cardamine.

### The dehydroxynocardamine BGC is expressed *in vivo*

We identified a 19 bp putative Diphtheria Toxin Repressor (DtxR) iron box operator sequence (5’-TTATGCAAGGTTTTCCTAT-3’) (76) situated upstream of *dnoB* (**Fig. 4A**). When iron is abundant, DtxR forms a complex with ferrous iron and binds iron box operator sequences to repress the transcription of downstream genes, but this repression is relieved when iron is scarce (77). Transcriptional profiling of *S. aureus* from the nasal cavity has indicated that bacteria colonizing the nose are iron-starved *in vivo* (78). Therefore, we wanted to determine if the dehydroxynocardamine BGC is expressed *in vivo.* As part of the human microbiome project (HMP) prediabetes study (63), nasal swabs from the anterior nares were collected from 16 individuals over multiple visits and processed for metatranscriptome sequencing. We analyzed the nasal metatranscriptomes for expression of the dehydroxynocardamine BGC that we identified and several housekeeping genes (*rpoB, gyrB, sigA*, and *rpsL*). We detected *dno* expression in 7/16 individuals from at least one visit (mean = 2.5 visits/subject, median = 1 visit/subject) with expression values ranging from 0.01 - 154.69 transcripts per kilobase million (TPM) (mean = 26.94 TPM, median = 3.44 TPM). For comparison, we detected expression of housekeeping genes at similar levels (**Fig. 5**), with the exception of *rpsL*, which is known to be more highly expressed than the other selected housekeeping genes in the model Gram-positive organism *B. subtilis* (79). Together, these results indicate that the dehydroxynocardamine BGC is expressed *in vivo.*

**Figure 5.**
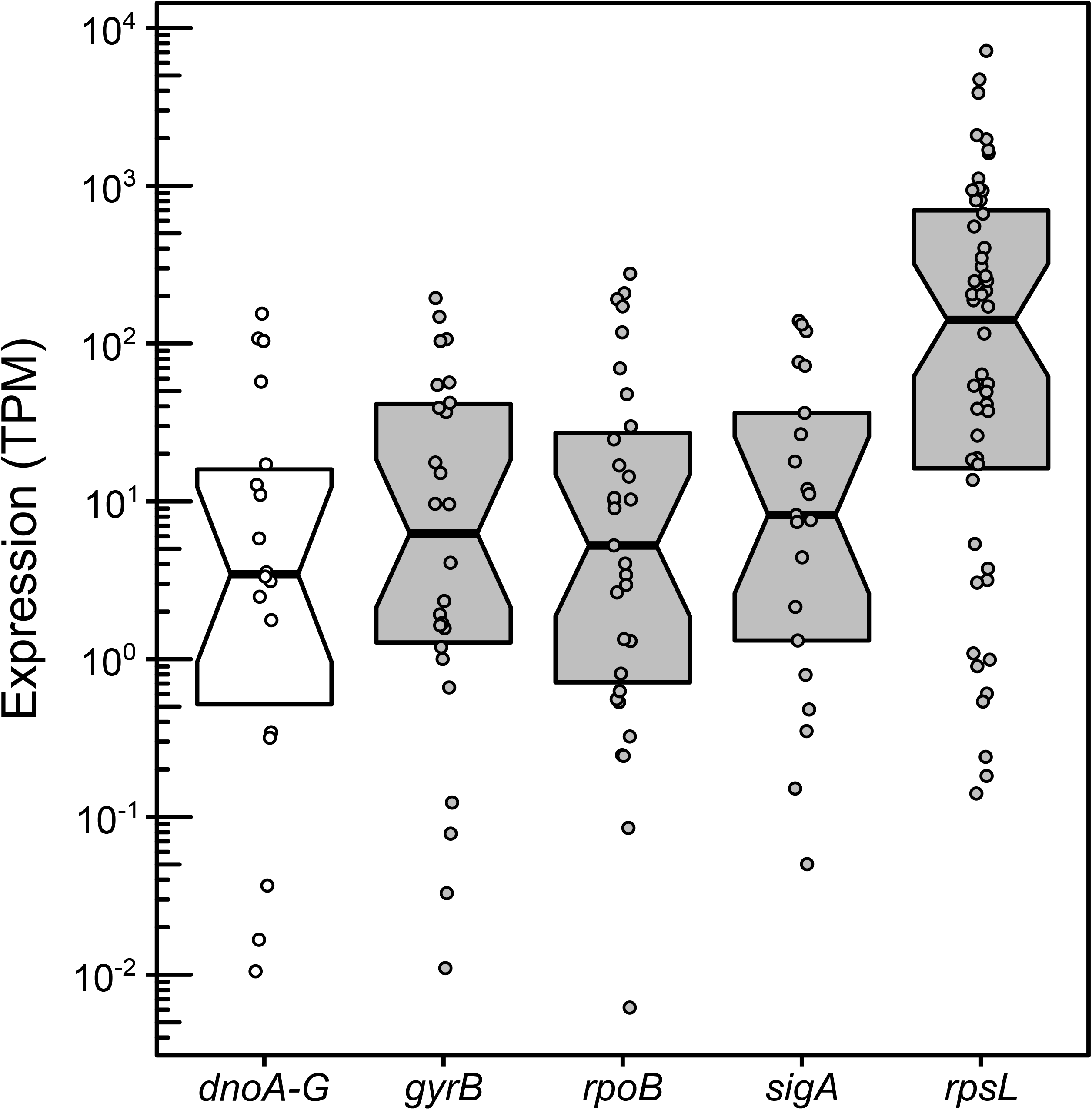
The dehydroxynocardamine BGC is expressed *in vivo.* 95 nasal metatranscriptomes from 16 subjects in the NIH HMP were pseudoaligned to the genome of *C. propinquum* HSID18034 to detect expression of the dehydroxynocardamine BGC (*donA-G*, white) and housekeeping genes *gyrB, rpoB, sigA*, and *rpsL* (gray). The upper and lower bounds of the notched box plot indicate the 75th and 25th percentiles, respectively. The bars indicate the median and the notches represent the 95% confidence interval of the median. Each dot indicates the expression of a single gene from a metatranscriptome in transcripts per kilobase million (TPM).

## DISCUSSION

In this study we investigated inhibition of nasal *Staphylococcus* spp. isolates by strains of sympatric Actinobacteria. We uncovered two distinct patterns of interactions: 1) *S. epidermidis* was significantly more sensitive than *S. aureus* to inhibition by nasal Actinobacteria, and 2) there was variability among *Corynebacterium* spp. in their ability to inhibit the growth of *S. epidermidis* (**Fig. 1**). Specifically, we noted that *C. propinquum* more strongly inhibited *S. epidermidis* when compared to inhibition by other *Corynebacterium* spp. Through a combination of comparative genomics (**Fig. 2**) and culture-based approaches (**Fig. 3**), we determined that the difference in inhibitory ability was due to production of a siderophore, dehydroxynocardamine (**Fig. 4**). By analyzing metatranscriptome samples from the anterior nares of human subjects, we established that the dehydroxynocardamine BGC is expressed *in vivo* (**Fig. 5**), indicating that iron-mediated exploitation competition is potentially relevant to competition and organismal fitness within the human nasal cavity.

One broad explanation for the difference in inhibition patterns between *S. aureus* and *S. epidermidis* strains (**Fig. 1B**) is that *S. epidermidis* and other coagulase-negative staphylococci (CoNS) lack the mechanisms that *S. aureus* uses to obtain iron from the hosts and their external environment. Strains of *S. epidermidis* and other CoNS do not grow well on iron-limited media (19) and, unlike *S. aureus*, are unable to scavenge iron from human transferrin (80). Our finding that iron supplementation rescues *S. epidermidis* from inhibition by *C. propinquum* (**Fig. 3B-C**) is consistent with previous results showing that *S. epidermidis* and other CoNS are sensitive to inhibition by desferrioxamine siderophores and other iron-chelating agents *in vitro* (81, 82). In contrast to *S. epidermidis*, strains of *S. aureus* may survive siderophore-mediated inhibition through production of their own carboxylate siderophores called staphyloferrin A (83) and B (84). Though the genomes of some strains of *S. epidermidis* and other CoNS encode BGCs for staphyloferrin, their siderophore activity is markedly decreased, indicating lower levels of siderophore production (85). As *S. epidermidis* strains lack the same mechanisms of iron acquisition and their siderophore production is under different regulatory control than *S. aureus* (85), they are susceptible to iron limitation imposed by *C. propinquum.* In addition, it has been previously reported that *Staphylococcus* spp. are able to use catecholate and hydroxamate siderophores produced by *Corynebacterium* spp. and other organisms (86). Perhaps in addition to higher levels of siderophore production, *S. aureus* is also better able to pirate dehydroxynocardamine produced by *C. propinquum* and circumvent inhibition.

Here, we note that siderophore-mediated inhibition does not fully explain the contrast between the results of interactions of *S. aureus* and *S. epidermidis* with nasal Actinobacteria. In addition to the three siderophore-producing strains of *Corynebacterium*, only five of the eleven other Actinobacteria encoded siderophore BGCs in their genomes (**Fig. 1B**). Therefore, these Actinobacteria likely use alternative mechanisms to inhibit the growth of *S. epidermidis*, such as interference competition mediated by antibiotic production, which is common among nasal bacteria (30–32). Subsequent work will be required to identify the molecules responsible for inhibition of *S. epidermidis* that are produced by non-siderophore producing nasal Actinobacteria.

Exploitation competition mediated by siderophore production is common among soil and marine bacteria (87–89), but within the context of host-associated systems, siderophore production is considered as a pathogen virulence factor (reviewed in 90) because mutants that are unable to produce siderophores are often defective in colonizing hosts and causing disease (91, 92). The view of siderophores solely as pathogen-produced molecules is strengthened by lipocalin-2, a protein component of the innate immune response that binds to catechol siderophores to prevent their uptake by bacteria and whose production is induced by bacterial colonization of mucosal surfaces (23, 24). However, thus far, little work has considered siderophore-mediated competition among the microbiota and between the microbiota and pathogenic bacteria (93). For instance, siderophore piracy is reported to occur between pathogens and beneficial or commensal bacteria in the human gastrointestinal tract (94, 95), but hitherto no such mechanisms have been reported within the human nasal cavity.

In conclusion, as 1) dehydroxynocardamine is not a known virulence factor, 2), *C. propinquum* is considered a normal part of the nasal microbiota (33, 96, 97), and 3) expression of *dnoA-G* was detected *in vivo* (**Fig. 5**), we propose that *C. propinquum* produces dehydroxynocardamine as a means to mediate exploitation competition for iron with other bacteria within the human nasal cavity.

## ACKNOWLEDGMENTS

We thank members of the Cameron Currie, James Gern, and Paul Straight (Texas A&M University) laboratories for feedback on this manuscript and project. We thank Aaron Stubbendieck for 3D printing stamps for rapid inoculation of co-culture inhibition assays. We thank Samuel McInturf for thoughtful discussion and assistance in R scripting. We thank Drs. Robert Lemanske Jr. and Daniel Jackson (University of Wisconsin-Madison), who are the principal investigators of the COAST birth cohort study, for providing nasal samples from which bacteria were isolated.

This work, including the efforts of Cameron R. Currie and James E. Gern, was funded by the NIH Centers for Excellence for Translational Research (U19-AI109673-01) and the NIH National Heart, Lung, and Blood Institute (P01 HL070831), respectively. The funders had no role in study design, data collection and interpretation, or the decision to submit the work for publication.

---

**Figure S1.**
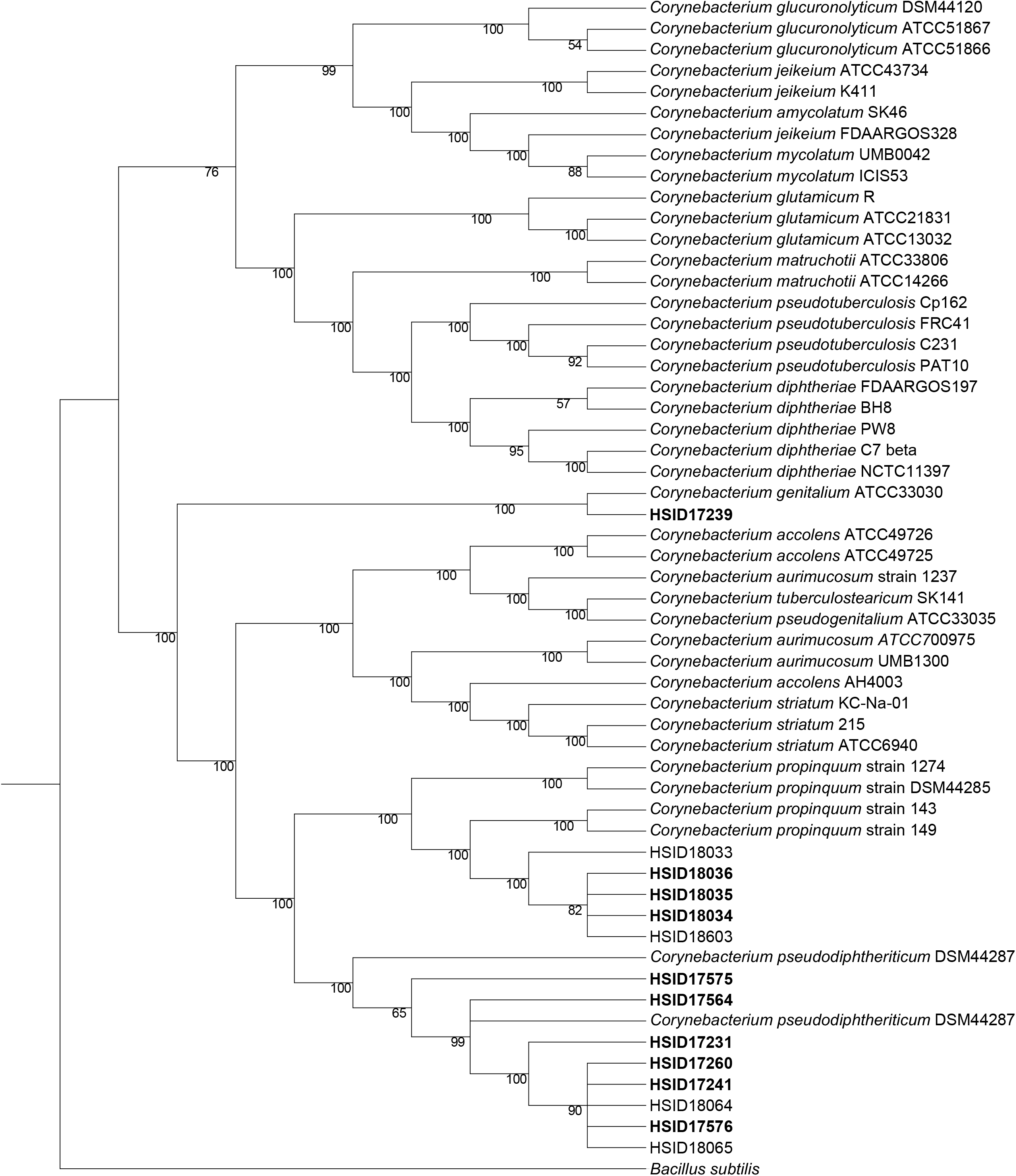
Core-genome cladogram of human-associated *Corynebacterium* spp. Core-genome cladogram of 55 human-associated *Corynebacterium* spp. built from 93 conserved, single copy genes. The strains of *Corynebacterium* spp. used in this study are in bold. The cladogram is rooted on *B. subtilis* 168 and nodes with less than 50% bootstrap support were collapsed. Nodes with ≥50% bootstrap support are indicated.

**Figure S2.**
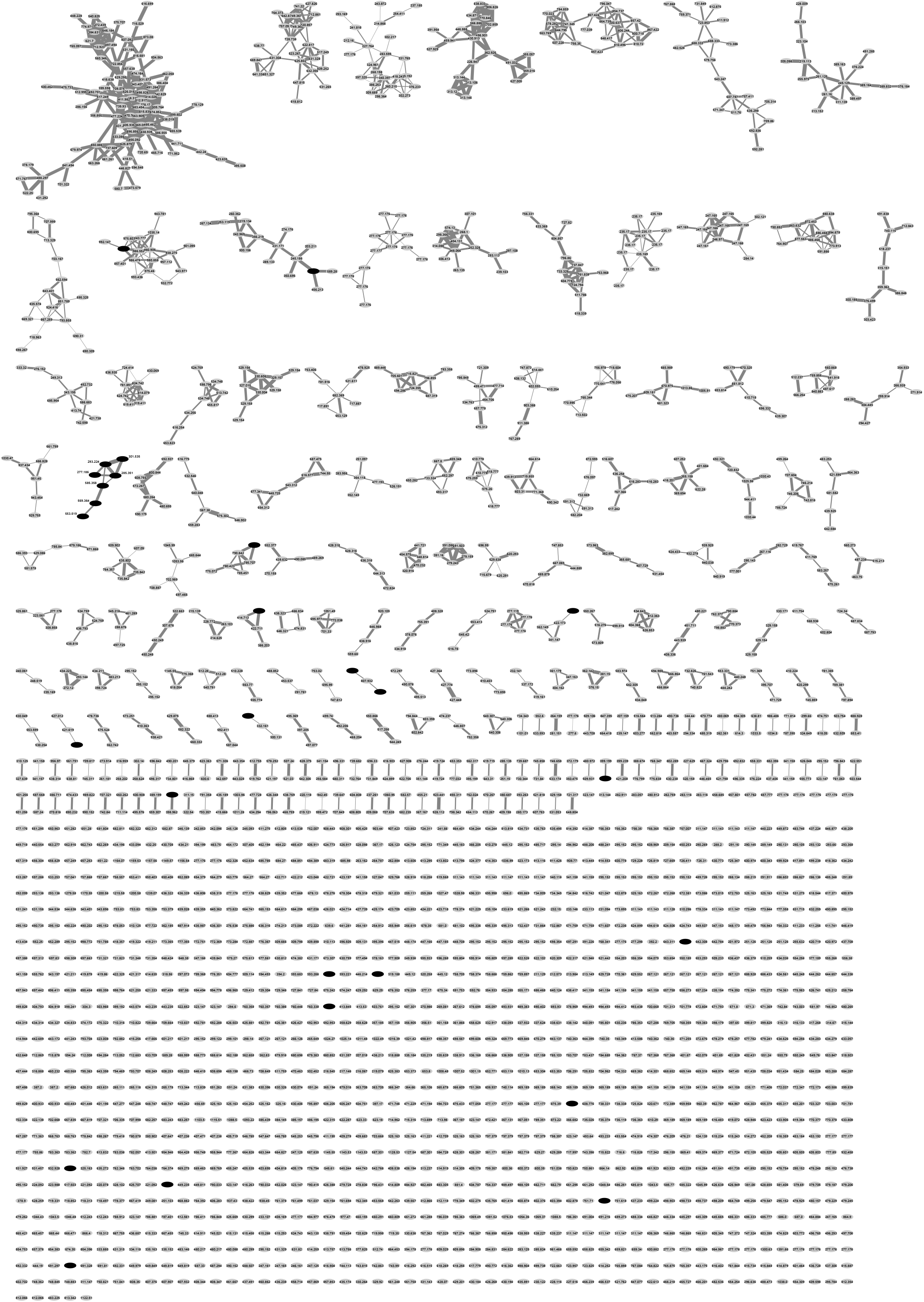
Full molecular network of *C. propinquum* and *C. genitalium* agar core extracts. Grey nodes are metabolites shared by both strains and black nodes are metabolites unique to *C. propinquum.* The edges are weighted to the cosine score between the two features.

**Figure S3.**
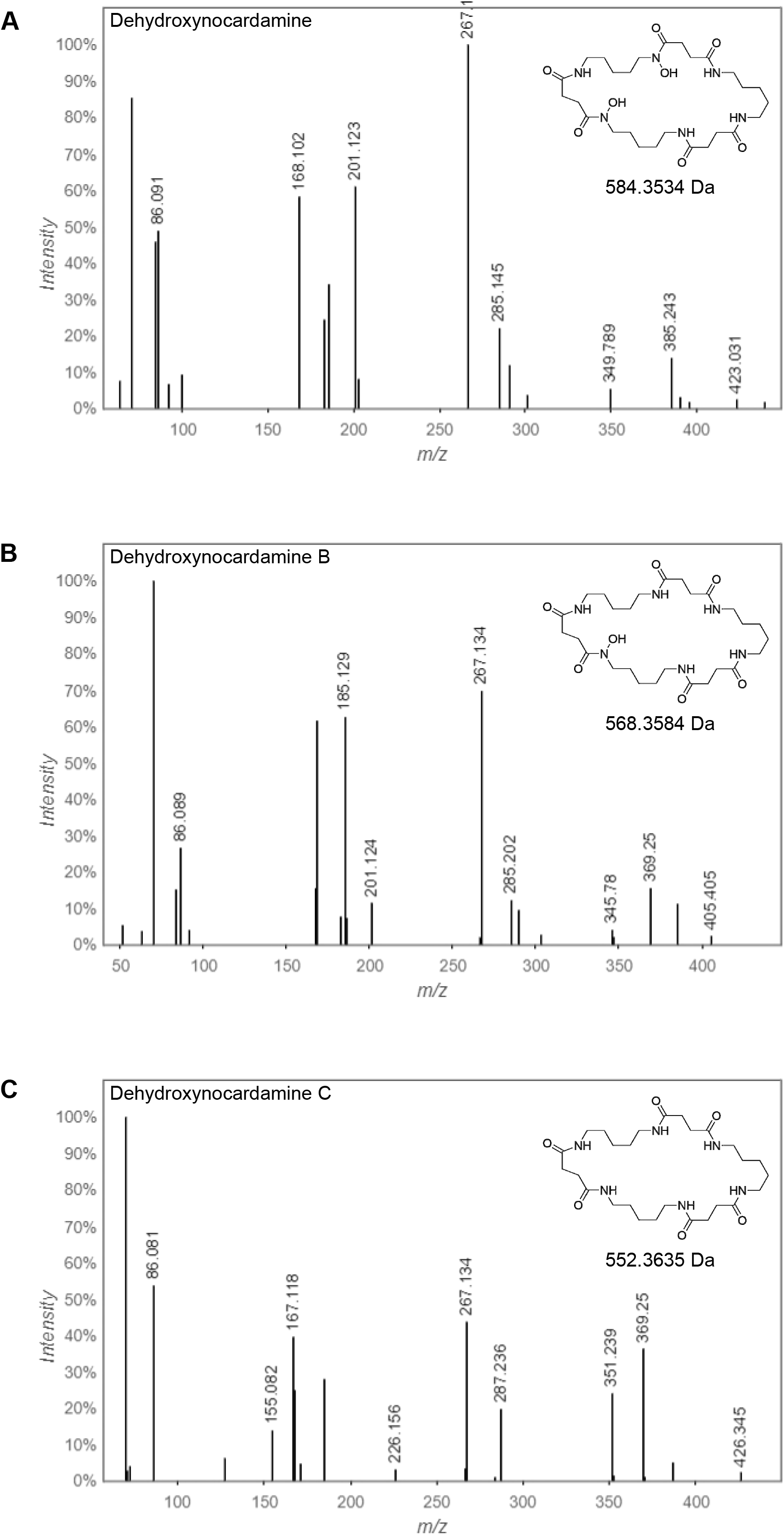
Tandem mass spectra of dehydroxynocardamine variants. The tandem mass spectra of **(A)** dehydroxynocardamine, **(B)**, dehydroxynocardamine B, and **(C)** dehydroxynocardamine C are shown. The insets show the structure of the dehydroxynocardamines and their molecular weights.

**Table S1.**
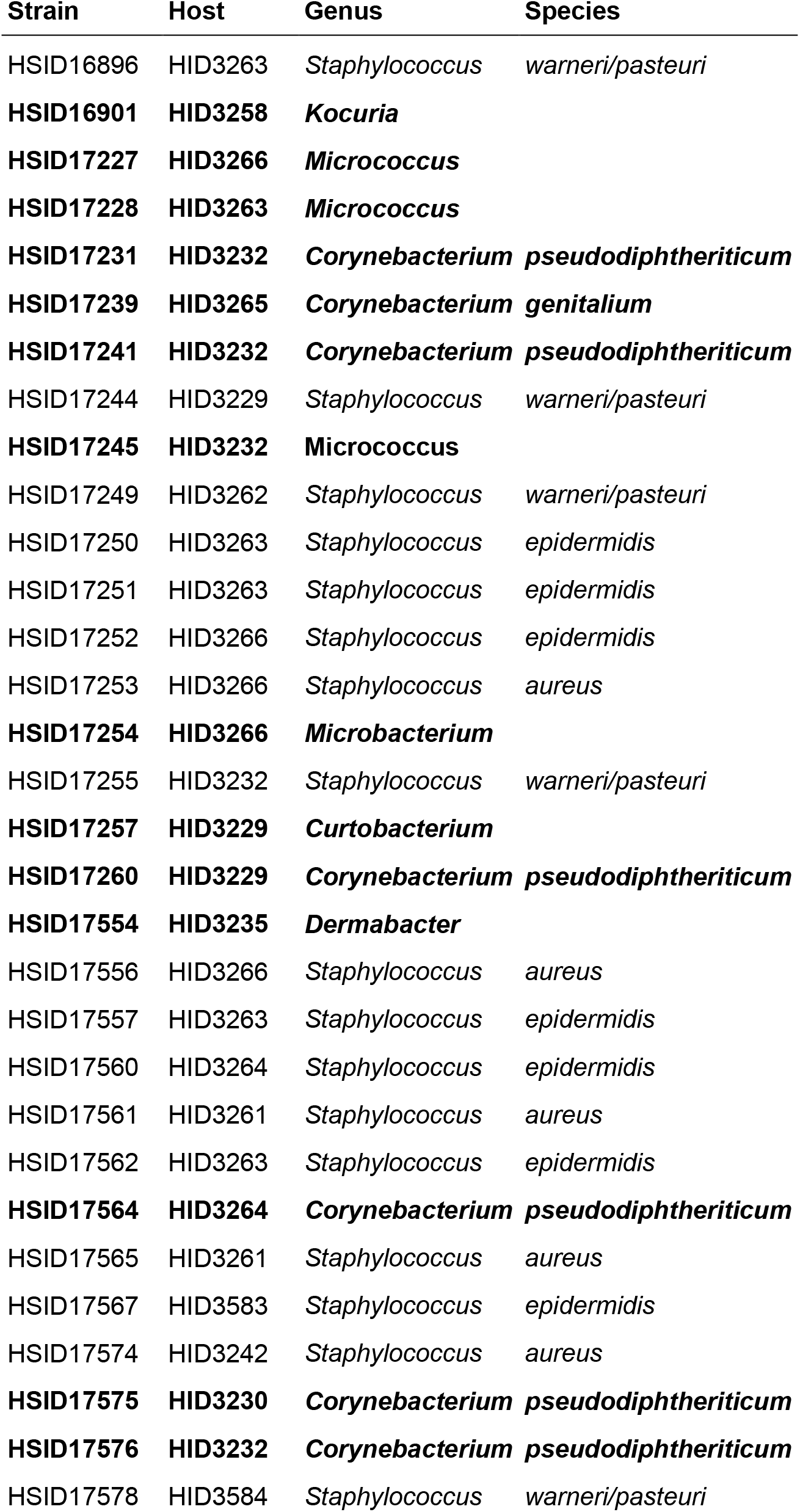
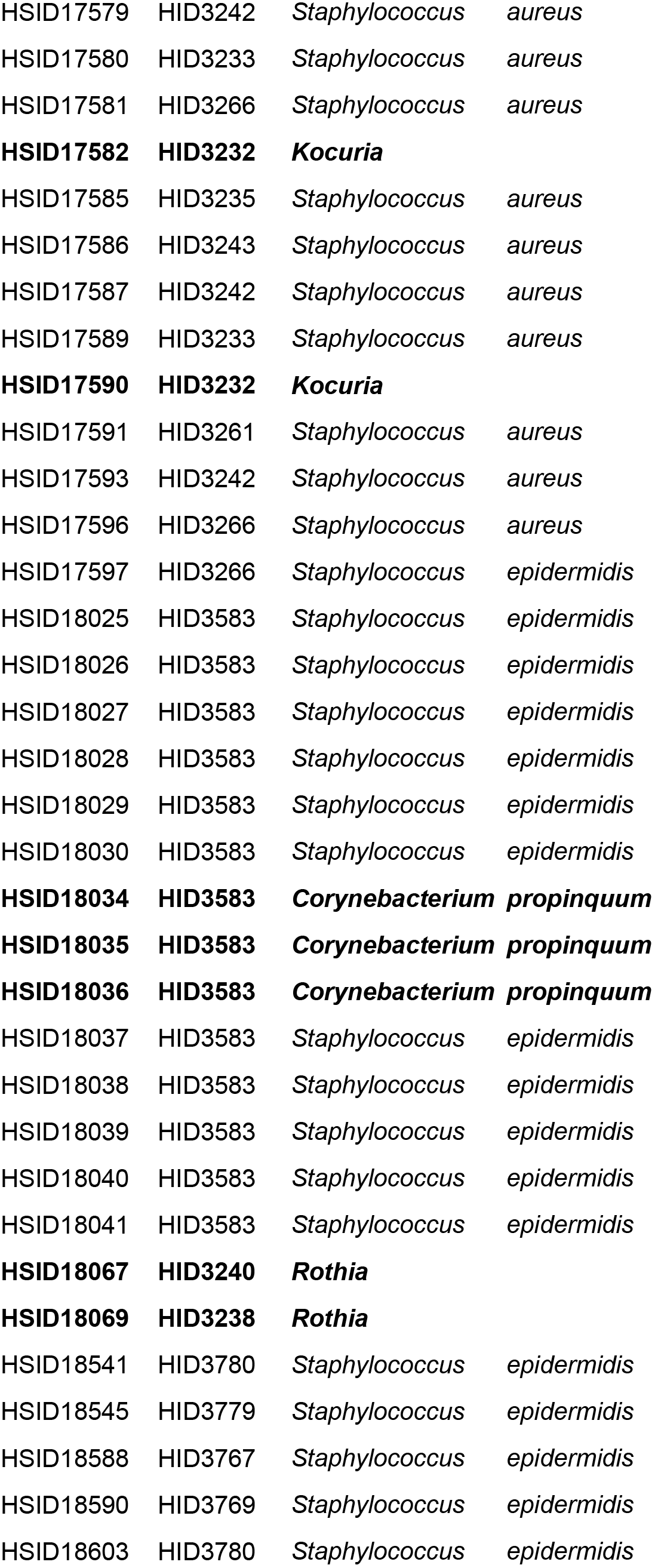
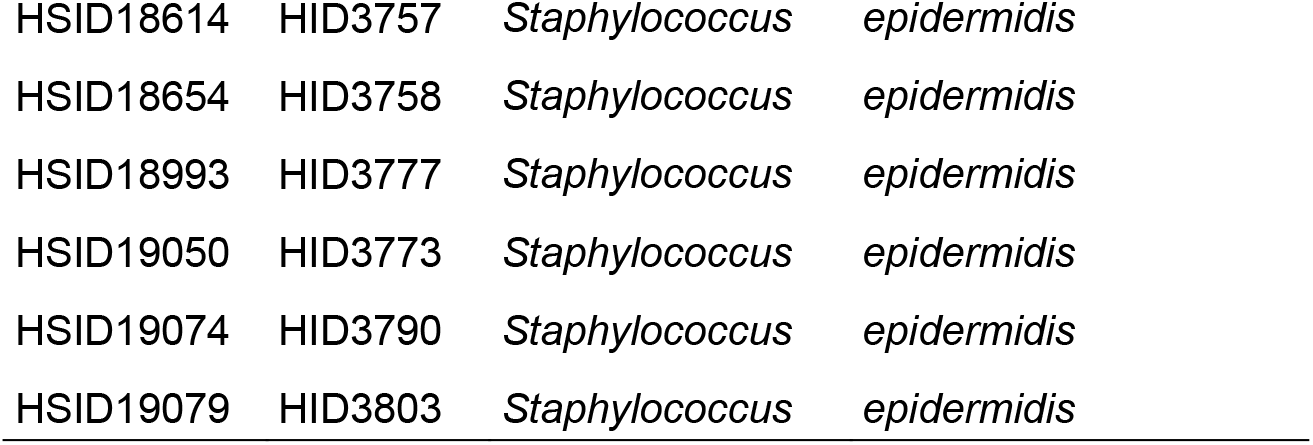
Strains of nasal Actinobacteria and *Staphylococcus* used in this study. Bolded strains indicate Actinobacteria with sequenced genomes.

